# OpenMebius2: GUI-based software for ^13^C-metabolic flux analysis with tracer labeling pattern suggestions for accurate flux predictions

**DOI:** 10.64898/2026.03.20.698926

**Authors:** Tatsumi Imada, Hiroshi Shimizu, Yoshihiro Toya

## Abstract

^13^C-metabolic flux analysis (^13^C-MFA) is a crucial technique that experimentally determines metabolic flux distribution. Although precision of each flux strongly depends on tracer labeling pattern, its optimization remains challenging. We developed an integrated platform, OpenMebius2, a graphical user interface (GUI)-based software for ^13^C-MFA that includes a tracer labeling pattern suggestion function to support subsequent experiments. The proposed function leverages metabolic flux distributions and their 95 % confidence intervals obtained using low-cost ^13^C-labeled substrates to evaluate hypothetical parallel labeling scenarios and predict improvements in flux estimation precision.

**Availability and implementation:** This software runs on Linux, macOS, and Windows. The source code and binary files are available at https://github.com/metabolic-engineering/OpenMebius2 under the PolyForm Noncommercial License 1.0.0.

## Introduction

Measurement of metabolic flux distribution, that is, the intracellular chemical reaction rate, is important to characterize the physiological state of a cell. Over the past decades, many technologies have been developed to measure intracellular metabolic flux distributions, among which ^13^C-metabolic flux analysis (^13^C-MFA) has emerged as a powerful and practical approach (Zamboni *et al*. 2009). ^13^C-MFA estimates the metabolic flux distributions from isotope labeling patterns of metabolites, based on ^13^C-labeling experiments. Accuracy of flux estimation is affected by multiple factors, including the pathway network structure, types of metabolites measured by mass spectrometry, degree of experimental errors, underlying flux distribution, and substrate ^13^C-labeling used in an experiment (Antoniewicz 2013). Experimental parameters must be optimized to estimate flux distributions with high accuracy.

For model microorganisms using glucose as a sole carbon source, the experimental protocols are well established for ^13^C-MFA. Previously reported studies provided experimental conditions and metabolic model information that can be easily utilized. However, appropriate ^13^C-labeling pattern of substrate may vary, even with minor modifications to the pathway network, such as introduction of heterologous pathways. Therefore, to estimate the flux with high accuracy, it is essential to examine suitable substrate labeling patterns for each condition. *In silico* simulations have often been performed to investigate optimal substrate ^13^C-labeling patterns (Metallo, Walther and Stephanopoulos 2009; Crown, Long and Antoniewicz 2016; Maeda *et al*. 2016; Toya, Ohashi and Shimizu 2018). To predict the optimal ^13^C-labeling of the substrate, metabolic flux distributions were randomly sampled, and corresponding mass distribution vectors (MDVs) of the metabolites were computed. Using the hypothetical experimental dataset, suitability of substrate ^13^C-labeling can be assessed by comparing the width of 95 % confidence intervals (CI) for the estimated flux. However, optimization of ^13^C-labeling patterns is computationally intensive and time-consuming, and its implementation requires a thorough understanding of ^13^C-MFA. Moreover, ^13^C-labeled substrates are often expensive, which can limit practical use of desired ^13^C-labeled substrates.

Many software packages related to ^13^C-MFA have been developed, such as 13CFLUX, Metran, INCA 2.0, OpenFLUX2, WUFlux, and mfapy, which offer high extensibility and rich functionality with a variety of features for optimizing substrate labeling (Weitzel *et al*. 2013; Shupletsov *et al*. 2014; He *et al*. 2016; Matsuda *et al*. 2021; Rahim *et al*. 2022; Stratmann *et al*. 2025). However, these software packages are primarily designed for users with advanced expertise and can be challenging for those who are unfamiliar with tracer experiments. Many researchers in the field of metabolic engineering who are not specialists in ^13^C-MFA only require determination of metabolic fluxes with sufficient accuracy for their specific research purposes. Here, we developed a function that suggests optimal labeling patterns for subsequent experiments tailored to end-user needs using metabolic flux distributions and their CI obtained from low-cost ^13^C-labeled carbon sources, and integrated it into OpenMebius, which has been used in our laboratory (Kajihata *et al*. 2014).

### The software

OpenMebius2 is a graphical user interface (GUI)-based software program implemented in MATLAB R2025a. Compiled binary versions of the software are available for Windows, macOS, and Linux, and it is freely accessible for non-commercial use. The source code and compiled binaries are hosted on GitHub (https://github.com/metabolic-engineering/OpenMebius2), with detailed documentation available online (https://metabolic-engineering.github.io/OpenMebius2/).

The new function, substrate labeling pattern suggestion for subsequent experiments, is based on the principle that parallel labeling experiments can improve the CI of estimated fluxes (Leighty and Antoniewicz 2013). Parallel labeling experiments involve performing independent culture experiments using substrates with different labeling patterns. By jointly analyzing multiple datasets obtained from these experiments, researchers can estimate a flux distribution with improved precision. While the potential precision of flux estimation by parallel labeling experiments can generally be assessed through substrate labeling optimization procedure in advance. In this software, we used CI obtained from existing experiments to further refine flux uncertainty. Specifically, assuming a parallel labeling scenario, tracer label optimization is performed to identify labeling patterns that would provide higher flux precision when required by a researcher. This approach enables researchers to evaluate both candidate labeled substrates for subsequent experiments as well as gain the expected benefit of parallel labeling in advance, thereby supporting informed decision-making in metabolic engineering studies.

An overview of the overall workflow is presented in Figure 1A. Researchers first perform culture experiments using low-cost labeled substrates, such as [1-^13^C]glucose or [U-^13^C]glucose, and obtain specific growth, substrate consumption, and product formation rates, and metabolite MDVs (STEP 1). Using the acquired dataset, a standard ^13^C-MFA is performed to determine the flux distribution and its CI (STEP 2). Based on the experimentally determined flux distribution, MDVs are computed for hypothetical additional labeled substrates. The CI are estimated using the MDVs obtained in STEP 1 together with the generated MDV dataset, assuming a parallel labeling experiment (STEP 3). Subsequently, the researchers design subsequent experiments by considering both the degree of improvement in flux precision and the cost of the required labeled substrates (STEP 4). By following these steps, researchers can obtain flux distributions with a level of precision tailored to their specific research goals.

**Figure 1.**
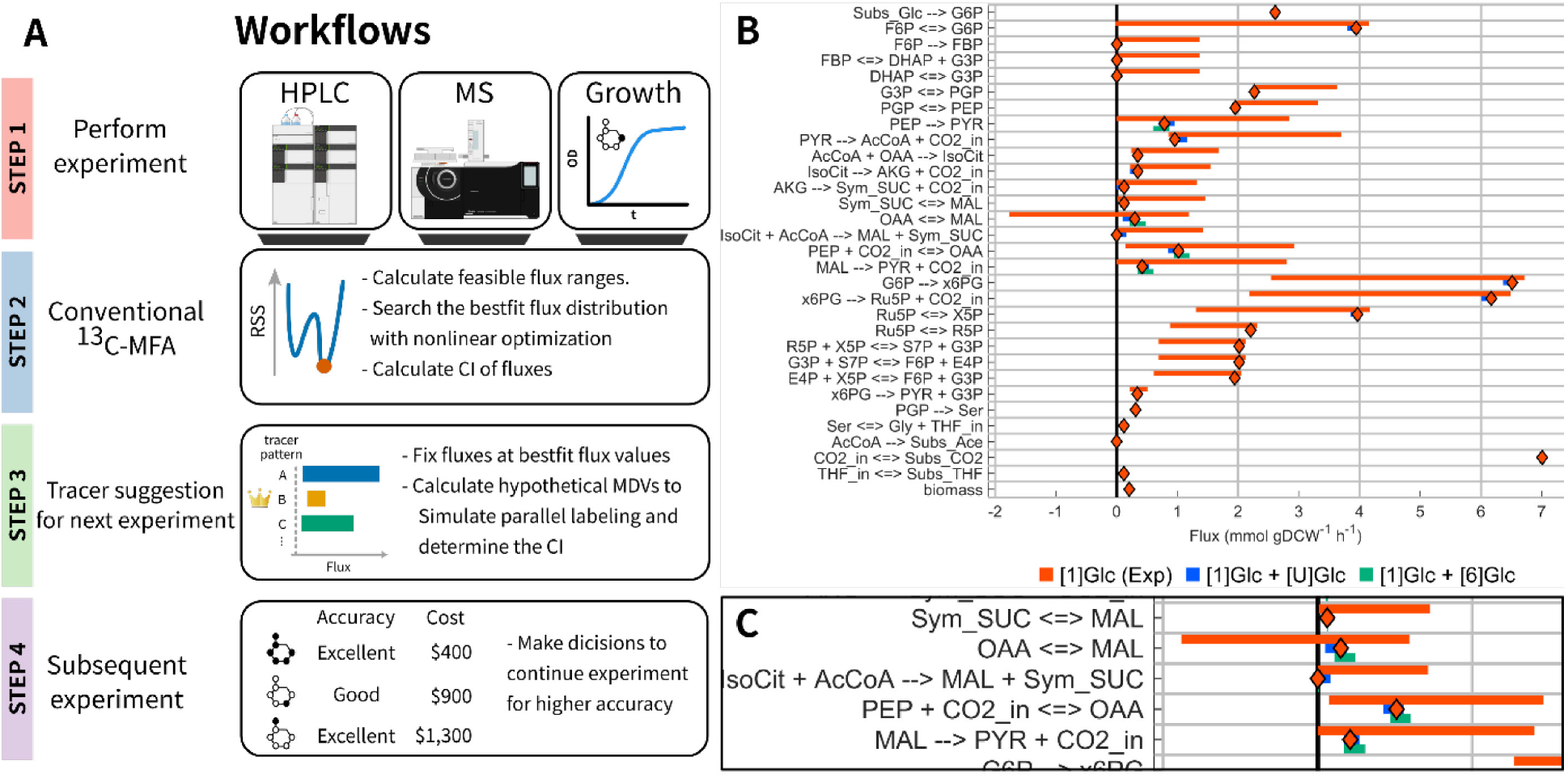
(A) Schematic workflows of ^13^C-MFA with tracer labeling pattern suggestion for subsequent experiments. Low-cost ^13^C-labeled substrates are first used to estimate metabolic flux distributions and their confidence intervals (CIs), which are then utilized to evaluate candidate tracers under hypothetical parallel labeling scenarios. (B) Example results of tracer labeling pattern suggestion. The red bar (top) indicates the CIs obtained from the original experimental dataset, while the blue (middle) and green (bottom) bars represent the predicted CIs assuming parallel labeling experiments with different tracer combinations. (C) Enlarged view of Figure 1B.

## Materials and Methods

A hypothetical flux distribution derived from *Escherichia coli* metabolism was generated, and both the model and dataset are available in the tutorial on the GitHub repository (https://github.com/metabolic-engineering/OpenMebius2). This model was based on the central carbon metabolism including glycolysis, the oxidative pentose phosphate pathway, tricarboxylic acid (TCA) cycle, Entner–Doudoroff pathway, and other anaplerotic pathways as shown in Table S1 and Table S2. To reflect a typical experimental setup for ^13^C-MFA, in which proteinogenic amino acid MDVs were obtained by mass spectrometry, MDVs were computed using an elementary metabolite unit-based model with a hypothetically generated flux distribution. Hypothetical MDVs of proteinogenic amino acid are shown in Table S3. This setup assumes the use of [1-^13^C]glucose, a common choice in ^13^C-MFA studies. Conventional ^13^C-MFA was then performed using this dataset to estimate the flux distribution and its CI. The optimal flux distribution was obtained by solving the following nonlinear optimization,

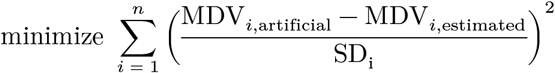

where *n* is the number of MDV element, MDV_*i*,artificial_ is the artificially generated MDV (generally this is an experimentally measured mass spectrum), MDV_*i*,estimated_ is the simulated MDV from the given metabolic flux distribution and MDVs of substrates, and SD_i_ is a standard deviation of each MDV element. In this study, we fixed SD_i_ is 0.01. CIs were calculated using a Monte Carlo method, in which CIs were estimated from the distributions of each flux obtained by adding random noise following a normal distribution to the original MDV values. To examine whether the accuracy of flux estimation could be further improved by using glucose ^13^C-labeled at a single carbon position, as well as other commonly used tracers, a substrate labeling suggestion was applied. The simulation was performed on a workstation equipped with a Xeon W7-3445 processor and 256GB of DDR5 ECC RAM, and took approximately 62 min to complete.

## Results and Discussion

Results of the example of substrate labeling are shown in Figure 1B. The red bar (top) in the plot represents CI obtained from the original experimental dataset. Blue (middle) and green (bottom) bars indicate the predicted CI, assuming parallel labeling experiments using [1-^13^C]glucose and [U-^13^C]glucose, or [1-^13^C]glucose and [6-^13^C]glucose respectively. In this example, the plot shows that the use of [U-^13^C]glucose is preferable to other substrates for subsequent tracer experiments to achieve a higher-precision metabolic flux estimation. In practice, researchers can make decisions by jointly considering the purpose of ^13^C-MFA in their research, cost of acquiring more accurate flux distributions, and expected improvement in accuracy achievable through additional experiments. We assessed cost-effectiveness by applying the incremental cost-effectiveness ratio (ICER), a metric commonly used in cost–effectiveness analyses (De Falco *et al*. 2022). ICER is defined as the additional cost required to achieve an incremental improvement in effectiveness. The improvement in flux estimation accuracy was quantified using a precision score (Crown, Long and Antoniewicz 2016). The CI of the fluxes obtained from the experimental data were used as the reference tracer, and maximum value of the precision score was set to 200. By definition, the precision score of the experimental dataset is equal to 1. Therefore, the ICER associated with performing an additional experiment *i* is defined as:

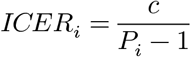

Where *c* denotes the cost of the additional labeled substrate ($ g^-1^), and *P*_*i*_ is the precision score calculated from simulation using tracer in a subsequent experiment *i*. The prices of the labeled substrates used in the simulations (as of December 2025) and their corresponding ICER values are summarized in Table 1. In this case study, although the use of [6-^13^C]glucose yielded the highest flux estimation precision, evaluation based on the ICER indicates that [U-^13^C]glucose provided the most favorable cost–effectiveness.

**Table 1.** Costs of labeled glucose tracers used in the simulations (as of December 2025) and their corresponding incremental cost-effectiveness ratio (ICER) values.

| Glucose tracers | Precision score | List price (\$ g-1) | ICER (\$ P-1) |
| --- | --- | --- | --- |
| [1-<sup>13</sup>C]glucose | 1 | 300 | - |
| [2-<sup>13</sup>C]glucose | 154 | 900 | 5.87 |
| [3-<sup>13</sup>C] glucose | 157 | 2200 | 14.11 |
| [4-<sup>13</sup>C] glucose | 163 | 2800 | 17.33 |
| [5-<sup>13</sup>C] glucose | 185 | 3500 | 19.02 |
| [6-<sup>13</sup>C] glucose | 186 | 1300 | 7.04 |
| [U-<sup>13</sup>C] glucose | 161 | 400 | 2.49 |
| [1,2-<sup>13</sup>C] glucose | 164 | 1200 | 7.38 |

## Conclusion

OpenMebius2 allows researchers to efficiently obtain high-precision metabolic flux distributions that are realistically achievable within practical experimental constraints. These workflows can be easily designed even by non-expert users, thereby providing an accessible entry point for researchers unfamiliar with ^13^C-MFA and for developers in industrial settings who wish to incorporate ^13^C-MFA as an evaluation method.

## Acknowledgements

We would like to thank Editage (www.editage.jp) for English language editing.

